# Electrophysiological signatures of visual recognition memory across all layers of mouse V1

**DOI:** 10.1101/2023.01.25.524429

**Authors:** Dustin J. Hayden, Peter S.B. Finnie, Aurore Thomazeau, Alyssa Y. Li, Samuel F. Cooke, Mark F. Bear

**Affiliations:** The Picower Institute for Learning and Memory, Department of Brain and Cognitive Sciences, Massachusetts Institute of Technology, Cambridge, MA, USA 02139; Kavli Institute for Systems Neuroscience and Centre for Neural Computation, Norwegian University of Science and Technology, Trondheim, Norway 7020; Department of Pharmacology and Toxicology, Temerty Faculty of Medicine, University of Toronto, Toronto, Ontario, Canada M5S 1A8; Brain Repair and Integrative Neuroscience Program, Centre for Research in Neuroscience, Departments of Medicine, and Neurology & Neurosurgery, The Research Institute of the McGill University Health Centre, Montreal General Hospital, Montreal, Quebec, Canada H3G 1A4; Biochemistry Program, Wellesley College, Wellesley, MA, USA 02481; Institute of Medical Science, Faculty of Medicine, University of Toronto, Toronto, Ontario, Canada M5S 1A8; The Medical Research Council Centre for Neurodevelopmental Disorders (MRC CNDD) and The Department of Basic and Clinical Neuroscience, Institute of Psychiatry, Psychology & Neuroscience, King’s College London, London, UK SE5 8AF

## Abstract

In mouse primary visual cortex (V1), familiar stimuli evoke significantly altered responses when compared to novel stimuli. This stimulus-selective response plasticity (SRP) was described originally as an increase in the magnitude of visual evoked potentials (VEPs) elicited in layer (L) 4 by familiar phase-reversing grating stimuli. SRP is dependent on NMDA receptors (NMDAR) and has been hypothesized to reflect potentiation of thalamocortical synapses in L4. However, recent evidence indicates that the synaptic modifications that manifest as SRP do not occur on L4 principal cells. To shed light on where and how SRP is induced and expressed, the present study had three related aims: (1) to confirm that NMDAR are required specifically in glutamatergic principal neurons of V1, (2) to investigate the consequences of deleting NMDAR specifically in L6, and (3) to use translaminar electrophysiological recordings to characterize SRP expression in different layers of V1. We find that knockout of NMDAR in L6 principal neurons disrupts SRP. Current-source density analysis of the VEP depth profile shows augmentation of short latency current sinks in layers 3, 4 and 6 in response to phase reversals of familiar stimuli. Multiunit recordings demonstrate that increased peak firing occurs to in response to phase reversals of familiar stimuli across all layers, but that activity between phase reversals is suppressed. Together, these data reveal important aspects of the underlying phenomenology of SRP and generate new hypotheses for the expression of experience-dependent plasticity in V1.

**Significance Statement:** Repeated exposure to stimuli that portend neither reward nor punishment leads to behavioral habituation, enabling organisms to dedicate attention to novel or otherwise significant features of the environment. The neural basis of this process, which is so often dysregulated in neurological and psychiatric disorders, remains poorly understood. Learning and memory of stimulus familiarity can be studied in mouse visual cortex by measuring electrophysiological responses to simple phase-reversing grating stimuli. The current study advances knowledge of this process by documenting changes in visual evoked potentials, neuronal spiking activity, and oscillations in the local field potentials across all layers of mouse visual cortex. In addition, we identify a key contribution of a specific population of neurons in layer 6 of visual cortex.

## Introduction

Passive exposure to innocuous sensory stimuli that do not reliably predict reward or punishment produces habituation of innate behavioral responses across a wide range of organisms (Thompson and Spencer, 1966; Rankin et al., 2009). This conservation of function reflects the fundamental importance of habituation, which putatively serves to reduce energy use and enable the allocation of attention towards salient elements of the environment. Habituation is accompanied by a range of effects on evoked neural activity in different model organisms (Grill-Spector et al., 2006; Cooke and Ramaswami, 2020), yet we do not yet have a clear picture of how this foundational form of learning is implemented within the mammalian brain. In a paradigm we have previously developed, head-restrained mice briefly exposed to an identical phase-reversing, oriented sinusoidal grating stimulus over successive days exhibit behavioral habituation that is accompanied by pronounced stimulus-selective response plasticity (SRP) in binocular primary visual cortex (V1). Specifically, visual-evoked potentials (VEPs) recorded in layer (L) 4 elicited by phase reversals of these progressively familiar stimuli undergo a significant potentiation across days. Multiple molecular manipulations of V1 disrupt both SRP and accompanying behavioral habituation to the familiar stimulus orientation, suggesting shared local mechanisms of synaptic modification (Cooke et al., 2015; Kaplan et al., 2016). Thus, SRP may offer a direct readout of plasticity in the early visual system contributing to long-term recognition memory and habituation processes. Although changes in primary sensory activity to passive and reinforced visual stimulation have been extensively characterized (Aton et al., 2014; Kato et al., 2015; Makino and Komiyama, 2015; Poort et al., 2015; Kaneko et al., 2017; Gao et al., 2021), the sites and essential mechanisms that yield the observed response patterns remain poorly defined. In the case of SRP, it is still unclear how plasticity manifests outside of superficial cortical layers. This information is critical to pinpoint the primary site(s) of synaptic modification that store this tractable and foundational form of memory.

Several striking attributes of SRP had previously suggested the occurrence of thalamocortical (TC) Hebbian synaptic plasticity. First, long-term potentiation (LTP) induced in V1 by theta-burst electrical stimulation of the dorsal lateral geniculate nucleus (dLGN) both mimics and occludes SRP recorded in V1 L4 (Cooke and Bear, 2010). Second, both LTP and SRP are dependent on NMDAR activity in V1 (Kirkwood and Bear, 1994; Collingridge, 2003; Frenkel et al., 2006; Cooke et al., 2015) and can be prevented by interfering with delivery of AMPARs to postsynaptic membranes (Frenkel et al., 2006). Third, SRP is eye-specific, indicating that modifications likely occur at TC synapses conveying information exclusively from one eye, prior to binocular integration by V1 neurons (Frenkel et al., 2006; Cooke et al., 2015). Hence, an early model proposed that long-term visual recognition memory is stored as NMDAR-dependent potentiation of TC synapses within V1 (Montgomery et al., 2021). Subsequent observations challenged this simple view, however. First, pharmacological elimination of intracortical activity that spares TC synaptic transmission in L4 abolishes SRP expression (Cooke and Bear, 2014; Kaplan et al., 2016). Second, the activity of specific populations of GABAergic inhibitory neurons in V1 is modified with visual experience and significantly diverges for familiar and novel stimuli (Hayden et al., 2021). Third, SRP expression in L4 VEPs is disrupted by selective perturbations of activity in cortical parvalbumin-expressing (PV+) interneurons (Kaplan et al., 2016). Fourth, selective knockout of NMDARs from principal cells in L4—the primary target of TC input to V1—spares both SRP of L4 VEPs and behavioral habituation (Fong et al., 2020). Fifth, short-latency responses recorded in L4 at the onset of familiar and novel stimuli are indistinguishable, with differences emerging only over hundreds of milliseconds and not manifesting in VEPs until the second phase reversal (Hayden et al., 2021). Together these findings suggest that SRP is not a direct readout of feedforward potentiation in L4. Instead, it is likely to involve intracortical plasticity and/or feedforward TC potentiation at still unidentified synapses outside of L4, which nevertheless influence responses in L4.

To advance our understanding of where and how synaptic plasticity in V1 serves visual recognition memory, in the current study we first investigated how manipulating NMDARs in neurons *outside* of L4 influences SRP measured with VEPs *within* L4, then went on to analyze SRP expression across all layers of V1. We confirmed that V1 glutamatergic neurons play a key role in SRP by using an intersectional genetic strategy to knock out NMDARs in only principal cells. We next targeted NMDARs in a genetically defined subset of excitatory neurons in L6 that are known to influence PV+ inhibition in L4 (Bortone et al., 2014; Guo et al., 2017), and found that the effect on SRP mimicked the pattern observed following NMDAR knockout in excitatory neurons across all layers of V1. The discovery that a molecular manipulation in the deep layers disrupts SRP manifesting in L4 VEPs motivated us to then use high-density linear electrode arrays to simultaneously record neural activity across all layers of V1 after 6 days of SRP induction. The observed changes in both superficial and deep cortical layers are consistent with data previously obtained from L4 (Cooke et al., 2015; Hayden et al., 2021). Namely, phase reversals of familiar stimuli evoke larger amplitude VEPs across all layers compared to novel stimuli. Blocks of familiar stimuli also increase low frequency oscillatory power across all layers, whereas novel stimuli increase high frequency power except in the deepest portions of visual cortex. We confirm the previous finding that there is elevated peak L4 unit activity immediately following each phase reversal of the familiar stimulus (Cooke et al., 2015) and extend this observation to reveal that this increase in peak firing rate is also apparent in superficial, middle, and deep layers. After this transient increase, however, there is an extended period of suppressed firing between phase reversals of familiar stimuli that is not apparent for novel stimuli. Thus, while the phasic spiking response produced by familiar stimulus onset is greater than for novel stimuli, the overall firing rate is reduced, in agreement with dominant repetition suppression literature (Grill-Spector et al., 2006) and previous calcium imaging studies (Kato et al., 2015; Makino and Komiyama, 2015; Kim et al., 2020). Together, these sustained changes during familiar innocuous stimuli are consistent with a shift in cortical processing mode that is influenced by NMDARs on L6 principal cells.

## Materials and Methods

### Animal subjects

All procedures adhered to the guidelines of the National Institutes of Health and were approved by the Committee on Animal Care at MIT, Cambridge, MA, USA. Mice were housed in groups of 2-5 same-sex littermates after weaning at P21-23. They had access to food and water *ad libitum* and were maintained on a 12-hour light-dark cycle. For acute laminar electrophysiological recordings, we used male and female C57BL/6J mice (The Jackson Laboratory, Bar Harbor, ME) bred in the MIT animal colony.

For the cell-type specific knock-out experiment, we used either an intersectional viral strategy in genetically modified mice or mouse lines were crossed to restrict *Grin1* deletion to subpopulations of excitatory neurons. In both cases, mice expressing loxP sites around both copies of the *Grin1* gene were used to ablate the GluN1 subunit and, thereby, functional NMDARs in a Cre-dependent manner (*Grin1*^fl/fl^, The Jackson Laboratory, RRID:IMSR_JAX:005246; (Tsien et al., 1996)). For the local excitatory cell-specific GluN1 knockout in V1, AAV8-CaMKIIa-Cre-GFP (knockout group) or AAV8-CaMKIIa-GFP (control group; UNC Vector Core, 4.4 × 10^11^ vg/mL in sterile saline) was injected bilaterally into binocular V1 of *Grin1*^fl/fl^ animals (see surgical methods, below). To target the GluN1 knockout to L6 cortico-thalamic neurons, *Grin1*^fl/fl^ mice were instead crossed with the previously described *Ntsr1-Cre* recombinase mouse line (B6.FVB(Cg)-Tg(Ntsr1-cre)GN220Gsat/Mmucd; *Ntsr1-Cre, Layer-6*, GENSAT, RRID:MMRRC_030648-UCD; (Gong et al., 2007)). Experimental subjects were *Grin1*^fl/fl^ and either Ntsr1-Cre^+/-^ (knockout group) or Ntsr1-Cre^−/-^ (control group). To histologically validate Cre-expression, *Ntsr1*-Cre mice were also crossed with a Cre-reporter mouse line (B6.Cg-Gt(ROSA)26Sortm14(CAG-tdTomato)Hze/J; Ai14-TdTomato, The Jackson Laboratory, RRID:IMSR_JAX:007914).

### Surgery

For layer 4 (L4) VEP recordings in cell-type specific knockout experiments (**Figures 1-2**), young adult mice were injected sub-cutaneously (s.c.) with 0.1 mg/kg Buprenex and 1.0 mg/kg Meloxicam to provide analgesia. Induction of anesthesia was achieved via inhalation of isoflurane (3% in oxygen) or intraperitoneal injection of ketamine (50 mg/kg) and xylazine (10 mg/kg). Thereafter, anesthetic plane was maintained via inhalant isoflurane (∼1-2% in oxygen) for the duration of surgery (30-60 mins). Prior to surgical incision, the head was shaved and the scalp cleaned with povidone–iodine (10% w/v) and ethanol (70% v/v). The scalp was resected along the midline and the skull surface was scored with a scalpel blade, and residual connective tissue removed with sterile saline and a cotton-tipped applicator. A steel headpost was affixed to the skull (anterior to bregma) with cyanoacrylate glue, which was used to position the mouse in a stereotaxic surgical frame (model 960 or 963, Kopf Instruments, Tujunga CA). Small burr holes were drilled above both hemispheres of binocular V1 (3.0-3.1 mm lateral of lambda). For *Grin1*^fl/fl^ mice receiving AAV injections, a glass capillary tube (model #3-000-203-G, Drummond Scientific Company, Broomall PA) was backfilled with mineral oil before being loaded into a Nanoject III injector (model #3-000-207, Drummond) affixed to the stereotaxic arm. The capillary was front-loaded with diluted AAV and gradually lowered ∼750 into V1, then retracted to 700 microns and permitted to equilibrate for 2-5 minutes before commencing injections. A total of 6 injection pulses of 13.8 nL were performed at each of 3 cortical depths (700, 450, and 250 microns), with 20 secs between pulses and 120 secs between depths. The pipette was slowly retracted 5-10 minutes after completion of the final injection, which reduces AAV backflow from the target site. For all NMDAR knockout mice, tapered blunt-tip 300-500 kΩ tungsten recording electrodes (model #30070, FHC, Bowdoin, ME, US; 125 μm diameter shank) were implanted in each hemisphere, 450 μm below cortical surface. Silver wire (A-M systems, Sequim, WA, US) reference electrodes were placed over right frontal cortex. Electrodes were secured using cyanoacrylate glue and the skull was covered with dental cement (Ortho-Jet, Lang Dental, Wheeling IL). Meloxicam (1 mg/kg) was administered daily upon return to the home cage for 48-72 hours following surgery. Signs of infection and discomfort were carefully monitored. Mice were allowed to recover for at least 48 hours prior to head-fixation.

**Figure 1 –.**
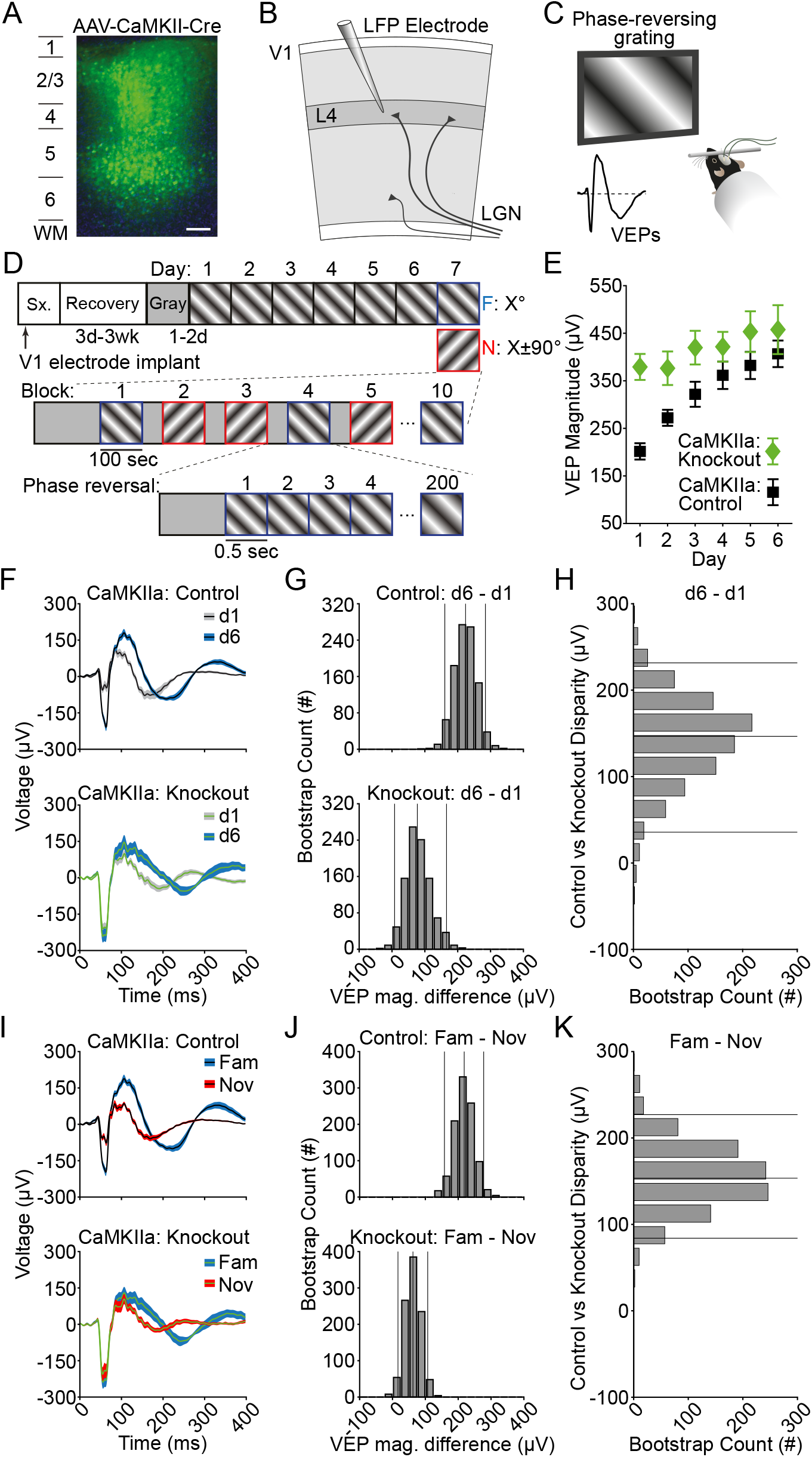
NMDA receptor knockout in V1 excitatory neurons affects L4 VEPs and SRP. (**A**) A representative confocal micrograph of virally-induced fluorescent protein co-expression following injection of AAV8-CaMKIIa-Cre-GFP – a proxy for the spatial distribution of Cre-mediated NMDA-receptor knockout across layers of V1 (green – GFP; blue – Hoechst counterstain; scale bar: 100 *µm*). (**B**) Diagram of visual cortex showing the electrode placement within L4. (**C**) Averaged VEPs elicited by phase reversals of full field, oriented, sinusoidal grating stimuli acquired binocularly from awake, head-fixed mice. (**D**) For 6 days, mice saw five blocks of phase-reversing gratings of a single orientation (100 phase reversal at 2 Hz), separated by 30 sec of gray screen. On day 7, five blocks of the now familiar orientation were pseudo-randomly interleaved with five blocks of a novel stimulus (rotated 90 degrees from the familiar orientation). (**E**) VEP magnitude plotted over days for both CaMKII-KO and control groups (symbol and bars represent mean ± SEM). (**F**) As a result of multiple days of experience, control animals had increased VEPs to the same stimulus on day 6 compared to day 1 (n = 13 mice). CaMKII-GluN1 KO animals did not show a similar increase (n = 14 mice). (**G**) The bootstrapped L4 VEP magnitude difference between day 6 and day 1 was large in control animals and small in KO animals. (**H**) The group disparity value is plotted, which compares bootstrapped day 6 – day 1 VEP magnitude differences between the control and knockout mice. The disparity shows that KO group undergoes less potentiation as a result of daily stimulus exposure. (**I, J, K**) Same as in (**F, G, H**), but comparing a novel stimulus on day 7 to the familiar stimulus on day 7. The control group has a larger VEP magnitude difference as a result of stimulus-dependent experience compared to knockout animals. The left and right vertical lines in **G** and **J** are the 95% confidence intervals generated by the bootstrap procedure. The middle vertical line is the median bootstrapped difference. Similarly, in **H** and **K**, the top and bottom vertical lines are the 95% confidence intervals for the disparity between the VEP magnitude differences of control and knockout animals, whereas the middle line is the median bootstrapped difference.

**Figure 2 –.**
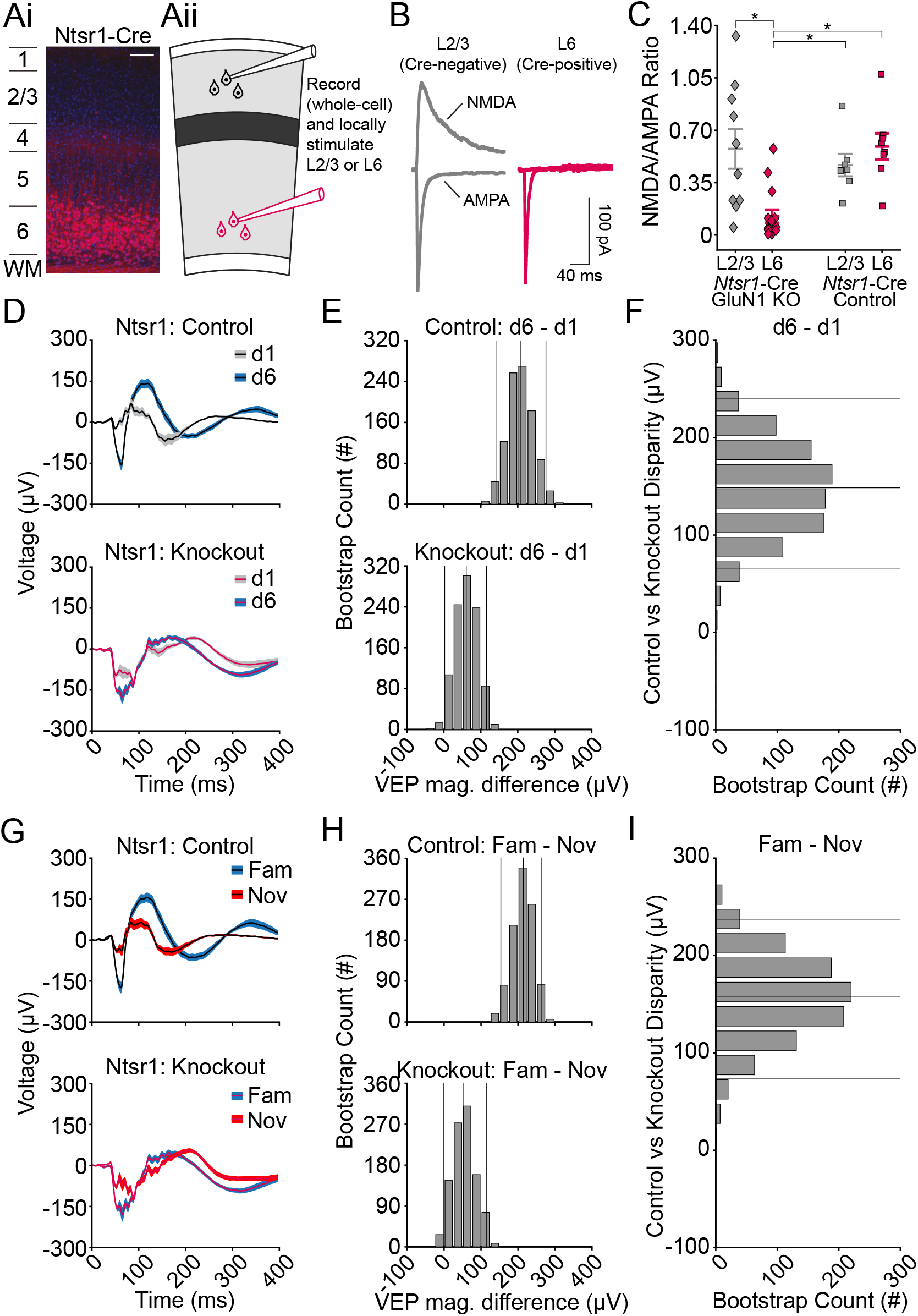
NMDA receptor knockout in L6 excitatory neurons affects L4 VEPs and SRP. (**Ai**) Representative confocal micrograph of fluorescent Cre reporter TdTomato across layers of V1 in a Ntsr1-Cre^+/-^, Ai14^+/-^ mouse, illustrating the pattern of NMDA-receptor knockout in Ntsr1-Cre^+/-^, Grin1^fl/fl^ mice (red – TdTomato; blue – Hoechst counterstain; scale bar: 100 *µm*). (**Aii**) Diagram of visual cortical slice showing electrode placement for L2/3 or L6 voltage clamp recordings during within-layer stimulation. (**B**) Sample traces of evoked NMDA and AMPA receptor currents from V1 principal cells in a L6-GluN1 knockout animal (Ntsr1-Cre^+/−^, GluN1^fl/fl^) recorded from a fluorescent (Cre-positive, right) and a non-fluorescent (Cre-negative, left) cell. Scale bars, 40 ms, 100 pA. (**C**) Mean NMDA/AMPA receptor current ratio from cells in animals possessing two floxed copies of the GluN1 allele (n = 4 animals, 10 cells in L2/3, 15 cells in L6) or from animals possessing at least one wildtype copy of GluN1 (n = 4 animals, 7 cells in L2/3, 8 cells in L6). Horizontal bars represent mean ± SEM. An asterisk indicates the compared cells exhibited a significant difference in NMDAR/AMPA current ratio, as determined by hierarchical bootstrapping. (**D**) As a result of multiple days of experience, control animals had increased VEPs to the same stimulus on day 6 compared to day 1 (n = 12 mice). Ntsr1-NMDAR KO animals showed a smaller increase over days (n = 15 mice). (**E**) The bootstrapped L4 VEP magnitude difference between day 6 and day 1 was observed in both control and KO animals. (**F**) The group disparity value is plotted, which compares bootstrapped day 6 – day 1 VEP magnitude differences between the control and knockout mice. The disparity shows that the L6 NMDA receptor knockout group undergoes reduced VEP magnitude change as a result of stimulus experience. (**G, H, I**) Same as in (**D, E, F**), but comparing a novel stimulus on day 7 to the familiar stimulus on day 7. The knockout group shows reduced levels of stimulus-dependent plasticity. The left and right vertical lines in **E** and **H** are the 95% confidence intervals generated by the bootstrap procedure. The middle vertical line is the median bootstrapped difference. Similarly, in **F** and **I**, the top and bottom vertical lines are the 95% confidence intervals for the disparity between the VEP magnitude differences of control and knockout animals, whereas the middle line is the median bootstrapped difference.

For laminar recordings, young adult C57BL/6J mice (n = 8 female, 10 male) were first injected with 5.0 mg/kg Meloxicam sub-cutaneously (s.c.) to provide analgesia. Induction of anesthesia was achieved via inhalation of isoflurane (3% in oxygen) and thereafter maintained via inhalant isoflurane (∼1-2% in oxygen). Prior to surgical incision, the head was shaved and the scalp cleaned with povidone–iodine (10% w/v) and ethanol (70% v/v). Scalp was resected and the skull surface was scored. Then, a stainless-steel head plate was attached to the skull and a 3 mm craniotomy was made over binocular V1. A sterile 3 mm round glass coverslip (CS-3R-0, Warner Instruments, Hamden, CT, USA) was gently laid on top of the exposed dura mater and held in place with VetBond (3M, St. Paul, MN) and thin Ortho-Jet bridges (Lang Dental, Wheeling, IL). A silver wire (A-M systems, Sequim, WA, US) reference electrode was inserted into left frontal cortex. Dental acrylic (C&B Metabond Quick Adhesive Cement System, Parkell, NY, USA) was mixed and applied throughout the exposed skull surface. The coverslip was covered with Kwik-Sil (World Precision Instruments, Sarasota, FL). Slow-releasing Buprenorphine was administered upon return to the home cage (1 mg/kg s.c.). Signs of infection and discomfort were carefully monitored. A total of 7 AAV-injected mice receiving were excluded from analysis due to poor recording quality, aberrant viral transduction/recombination, and/or health/technical issues.

On the morning of the acute recording, mice were once again isoflurane anesthetized and injected with 0.1 mg/kg Buprenex sub-cutaneously (s.c.) to provide analgesia. The Kwik-Sil was removed, the Ortho-Jet bridges broken, and the coverslip carefully pried off with a hooked needle. The craniotomy and surrounding area were filled with HEPES + agar. This mixture consists of 0.2 g high EEO Agarose (Sigma-Aldrich), 0.21 g Certified Low-Melt Agarose (Bio Rad), and 10.25 mL ACSF + HEPES (150.0 mM NaCl, 2.5 mM KCl, 10.0 mM HEPES, 2.0 mM CaCl2, 1.0 mM MgCl2). Kwik-Sil was applied again to prevent the agar from drying. This process was typically completed within an hour. Nonsteroidal anti-inflammatory drugs were administered upon return to the home cage (meloxicam, 1 mg/kg s.c.). The mice were head fixed, the Kwik-Cast removed, and a 64-channel laminar probe (Cambridge NeuroTech, Cambridge, England) was inserted slowly (∼100μm/minute) into V1 perpendicular to the cortical surface. Recordings were obtained and the mouse was euthanized immediately thereafter. We eliminated six mice due to either poor electrode placement (as measured by histology or non-stereotypical VEPs, n = 3), user error in recording (n = 1), superficial damage (n = 1), or surgical loss (n = 1). The remaining animals (n = 12; 6 female, 6 male) were included in all laminar analyses.

### Acute slice whole-cell electrophysiology

Functional loss of NMDA receptor expression in *Ntsr1-Cre* x *Grin1*^fl/fl^ animals was confirmed via *ex vivo* electrophysiology in slices obtained from dedicated cohorts of experimentally naïve mice. Fluorescence-guided whole-cell voltage clamp recordings were used to measure AMPA and NMDA receptor mediated excitatory post-synaptic currents (EPSCs) in Cre positive-cells from knockout mice (Cre^+/-^, *Grin1*^fl/fl^) and age-matched control animals hemizygous for Cre but with at least one wildtype copy of *Grin1* (i.e. Cre^+/-^, *Grin1*^fl/+^ or Cre^+/-^, *Grin1*^+/+^). Two strategies were used to fluorescently label cells with Cre recombinase activity. First, we injected an adeno-associated virus (AAV5-EF1α-DIO-GFP, UNC viral core) into the binocular zone of V1 to drive Cre-mediated expression of the green fluorescence protein (GFP) reporter (3.1 mm lateral of lambda, 81 nL of virus at each of 3 depths: 600, 450, and 300 μm from the cortical surface), allowing 3-4 weeks recovery prior to tissue harvest. Surgical injections were performed using a Nanoject II system (model #3-000-204, Drummond Sci.) as described above, except the mice were positioned in the stereotaxic frame using earbars. Second, we bred a triple transgenic animal using the Cre-driver x *Grin1*^*f*l/fl^ lines crossed with the Cre-dependent tdTomato reporter line, Ai14. In both cases, animals were approximately 6 months old at time of slice preparation. Coronal slices of V1 were prepared at a thickness of 350 μm in ice-cold dissection buffer containing (in mM): 87 NaCl, 75 Sucrose, 2.5 KCl, 1.25 NaH2PO4, 25 NaHCO3, 0.5 CaCl2, 7 MgSO4, 1.3 ascorbic acid, and 10 D-glucose, saturated with 95% O2 and 5% CO2. Slices were recovered for 40 minutes at 33°C and for approximately 1 hour at room temperature in artificial cerebrospinal fluid (aCSF) containing (in mM): 124 NaCl, 5 KCl, 1.23 NaH2PO4, 26 NaHCO3, 2 CaCl2, 2 MgCl2, and 10 D-glucose, saturated with 95% O2 and 5% CO2. Whole-cell patch clamp recordings were performed in continuous perfusion of carbogenated artificial cerebrospinal fluid (aCSF) at 30 °C using borosilicate pipettes with tip resistances of 3-5 MΩ. Pipettes were filled with balanced intracellular solutions containing (in mM): 115 cesium methane-sulfonate (CsMeSO3), 2.8 NaCl, 0.4 EGTA, 4 ATP-Mg2+, 10 Na+-phosphocreatine, 0.5 Na+-GTP, 5 TEA-Cl-, 5 QX-314 Br-buffered with 20 HEPES (pH 7.25, 290 mOsm). Layer 6 and layer 2/3 EPSCs were evoked by stimulation of layer 6 and layer 2/3, respectively, in the *Ntrs1*-Cre line (150 μs, 0.1 Hz, glass pipette electrode, and WPI A365 stimulus isolator) at holding potentials of -70 and +40 mV. The AMPA receptor component was measured from evoked EPSCs at -70 mV in the presence of picrotoxin (100 μM) and glycine (1 μM), and the NMDA receptor component was measured from evoked EPSCs at +40 mV in the presence of DNQX (20 μM). NMDA/AMPA receptor mediated EPSC ratios were calculated on a cell-by-cell basis and hierarchical bootstrapping was used to evaluate differences between groups.

### Visual stimulus delivery

For the cell-specific knockout mouse lines, after recovery from electrode implantation, experimentally naïve mice were acclimated to head restraint in front of a gray screen for a 30-minute session on each of two consecutive days. After this acclimation period, mice received 6 days of passive exposure to an oriented grating stimulus at a set, non-cardinal orientation. Each session began with 5 minutes of gray screen, after which they were presented with 5 blocks of 100 phase-reversals of an oriented grating stimulus, phase-reversing at 2 Hz. A 30-second period of gray screen was presented between each stimulus block. On day 7, mice were shown blocks of the familiar stimulus orientation pseudorandomly interleaved with blocks of a novel stimulus offset by ±90°.

For the laminar recordings, after recovery from the headplate surgery, mice were acclimated to head restraint in front of a gray screen for a 60-minute session on each of two consecutive days. After habituation, mice were presented with 10 blocks of 100 phase-reversals of an oriented grating stimulus phase-reversing at 1 Hz. They were shown this stimulus for 4 consecutive days. On day 5, they were shown five blocks of the familiar stimulus orientation pseudo-randomly interleaved with five blocks of a novel stimulus offset 90° from the familiar orientation. Each stimulus block was preceded by a period of gray screen, a period of black screen, and another period of gray screen. Gray periods and black periods lasted 10 seconds each, for a total of 30 seconds of pre-block activity. Discrete sections of gray and black screen viewing were timestamped for later normalization.

Visual stimuli consisted of full-field, 0.5 cycles/degree, 100% contrast, sinusoidal gratings that were presented on a computer monitor. Visual stimuli were generated using custom software written in either C++ for interaction with a VSG2/2 card (Cambridge Research systems, Kent, U.K.) or Matlab (MathWorks, Natick, MA, U.S.) using the PsychToolbox extension (http://psychtoolbox.org) to control stimulus drawing and timing (https://github.com/jeffgavornik/VEPStimulusSuite). Grating stimuli spanned the full range of monitor display values between black and white, with gamma-correction to ensure gray-screen and patterned stimulus conditions are isoluminant.

### In vivo electrophysiology experimental design and analysis

Electrophysiological recordings were conducted in awake, head-restrained mice. Recordings were amplified and digitized using the Recorder-64 system (Plexon Inc., Dallas, TX, US) or the RHD Recording system (Intan Technologies, Los Angeles, CA, US). For the cell-specific knockout mice, two recording channels were dedicated to recording continuous local field potential from V1 in each implanted hemisphere. Local field potential was recorded from V1 with 1 kHz sampling using a 500 Hz low-pass filter. For the laminar recordings on the Intan system, we sampled at 25 kHz and used a 0.1 Hz high pass and a 7.5 kHz low pass filter. Local field potential data was imported (see Importing and data cleaning) and the local field potential’s spectral content was analyzed (see Spectral analysis).

### Data import and cleaning

All analyses were conducted using custom MATLAB code and the Chronux toolbox (Bokil et al., 2010). For cell-specific knockout mice, the local field potential data was imported and converted to microvolts (µV). A handful of these recordings had minor errors in the event codes that were corrected post-hoc. For laminar recordings, the raw 25 kHz data from each channel was extracted and converted to µV. Then it was downsampled to 1000 Hz and a 3rd order 1-300 Hz Butterworth filter was applied. For all data, the mean of the entire channel’s data was subtracted from each timepoint to account for any DC offset in the system. Next, the data was locally detrended using the locdetrend function of the Chronux toolbox using a 0.5 s window sliding in chunks of 0.1 s. Finally, a 3rd-order Butterworth filter was used to notch frequencies between 58 and 62 Hz. For the multiunit activity of laminar recordings, the raw 25 kHz data was extracted for each channel. A 60 Hz, 10 dB bandwidth IIR notch filter was applied to each channel and the median value of each channel was subtracted from said entire channel. Finally, the median value across channels for each timepoint was subtracted from all channel’s timepoints. These data were stored separately for later use (see Multiunit activity analysis). Visually evoked potentials were normalized by subtracting the average of the first 10 milliseconds of each trial from that trial.

### Current-source density analysis and laminar identification

Performing a current-source density (CSD) analysis on laminar local field potential (LFP) data in V1 allowed us to identify layer 4 (L4) and align our data (Mitzdorf, 1985, 1987; Aizenman et al., 1996). The LFP data was temporally smoothed with a 20 ms Gaussian window. We then used a 5-point hamming window to compute the CSD using the standard formula (Ulbert et al., 2001; Speed et al., 2019). We refer to L4 as 0 µm - the site of the earliest and deepest sink immediately below the superficial source. All other channels were referenced according to that landmark. Thus, superficial layers were the channels above 0 µm and deep layers were the channels below 0 µm. For the sake of reporting in the **Results** section, we have broken the laminar data into four segments: L2/3 is above +90 µm, L4 is between +90 and -70 µm, L5 is between -70 and -210 µm, and L6 is below -210 µm. Reported statistics look for the largest significant difference within those four bounds.

### Spectral analysis

Given that the visually evoked potential violates assumptions required for spectral analysis (namely second-order stationarity), we only analyzed the spectral activity between 400 ms and 1000 ms after a phase-reversal. We computed the multi-tapered spectrum of the local field potential using the Chronux toolbox (Bokil et al., 2010). We used zero-padding to the 2nd power, 5 tapers, and a time-bandwidth product of 3. To calculate the normalized spectrum/spectrogram, we found the median spectrum/spectrogram of the animal’s black screen and took 10*log10(stimulus_spectrum/median_black_spectrum). This is reported as a decibel (dB).

### Multiunit activity analysis

To obtain the multiunit activity, we calculated a value known as multiunit activity envelope (MUAe). MUAe provides an instantaneous measure of the number and size of action potentials of neurons in the vicinity of the electrode tip (Legatt et al., 1980; Brosch et al., 1995; Super and Roelfsema, 2005). It does not depend on the arbitrary positioning of a threshold level and therefore does not select only large spikes. To obtain this value, a 3rd order bandpass (500 – 5000 Hz) Butterworth filter was applied to the common median referenced data that was previously stored (see Importing and data cleaning). Next, the absolute value of the data was taken (units of µV). Then a 3rd order low pass (< 250 Hz) Butterworth filter was applied and the data was downsampled to 1000 Hz. For z-scored data, the average across all trials for a given stimulus was calculated. Then, for each channel and animal, the average and standard deviation of the familiar and novel MUAe post-phase-reversal (up to 400 ms) was found. Both the familiar and novel mean and standard deviation were averaged together such that both familiar and novel stimuli could be z-scored to the same values (thus allowing direct comparison). To complete the z-score, the familiar MUAe data and the novel MUAe data were z-scored using the same averaged mean and standard deviation.

### Statistics

All statistics were done with the non-parametric hierarchical bootstrap for multi-level data (Saravanan et al., 2020). Briefly, statistical comparisons were between two groups, designated A and B. Most of our experiments utilize a within-animal design wherein animals experienced both familiar and novel stimuli (or stimuli on day 6 and day 1). In these cases, each animal in the experiment can and will contribute to both Group A and Group B. However, when comparing one genotype to another, each mouse can only contribute to the one appropriate genotypic group. To begin the bootstrap process, mice were randomly selected with replacement from the experimental population. For within-animal design comparisons, a random selection (with replacement) of mice was chosen and for each randomly chosen mouse a random selection (with replacement) of both Group A trials and Group B trials were selected. For genotype comparisons, a random selection (with replacement) of mice was chosen from each genotype and for each randomly chosen mouse a random selection (with replacement) of trials were selected to go into either Group A or Group B based on the genotype. Once all data were randomly selected, a statistic (e.g., VEP magnitude, spectral power, multiunit activity, etc) was computed from the trial and the mean difference between group A and group B was stored. This entire bootstrap process was repeated 1000 times. Once all 1000 bootstraps had been completed, the bootstrapped differences were sorted from lowest to highest value. The 500th value was the median group difference and is plotted in most graphs. The 25th value was the lower bound of the 95% confidence interval and the 975th value was the upper bound of the 95% confidence interval. Since the acute laminar recording experiments made multiple comparisons, we used the more conservative 99% confidence interval when making inferences. These used the 5th and 995th values of the sorted bootstrap. If the 95% or 99% confidence interval did not include zero, we report a statistically significant difference between group A and group B. This is reported as either an asterisk near the corresponding data on the plot or as a separate three-color plot with the valence and significance corresponding to a given color.

## Results

### Deletion of NMDARs in V1 principal cells disrupts SRP

In canonical models of the neocortical microcircuit, excitatory cells in layers 4 and 6 receive the bulk of feedforward projections from sensory thalamus (Douglas and Martin, 2004). Yet, surprisingly, selective knockout of NMDARs from the excitatory cells in L4 has no impact on SRP (Fong et al., 2020). Given the critical involvement of PV+ cells in SRP expression (Kaplan et al., 2016), the differential modulation of distinct V1 interneuron populations by familiar and novel stimuli (Hayden et al., 2021), and the fact that PV+ neurons receive excitatory input from the thalamus (Cruikshank et al., 2007), we first sought to confirm whether SRP relies on NMDARs on excitatory cells at all.

To do so, we knocked out the obligatory GluN1 subunit of the NMDAR in excitatory neurons of V1 by taking advantage of their selective expression of alpha Calcium/Calmodulin-dependent Kinase II (CaMKIIa). Recombination-mediated excision of the *Grin1* gene was targeted to excitatory cells across all cortical layers by intracranial microinjection of AAV8-CaMKIIa-Cre-GFP into transgenic mice in which Cre-dependent loxP sites were inserted around both copies of the *Grin1* gene (*Grin1*^fl/fl^). Thus, NMDARs were functionally ablated in only V1 excitatory neurons, as illustrated by histological examination of GFP expression induced by the AAV (**Figure 1A**). Control mice were infused with AAV8-CaMKIIa-GFP to produce equivalent fluorophore expression in the same excitatory cell population without Cre-mediated ablation of NMDARs. During the same surgery, local field potential recording electrodes were implanted within layer 4 and headposts affixed to the skull surface of all mice for subsequent acquisition of local field potential (LFP) data from V1 (**Figure 1B**). Following 3-4 weeks of surgical recovery and 2 days acclimation to the experimental apparatus, awake, head-fixed mice viewed 5 minutes of gray screen, before the onset of isoluminant full field, 2 Hz phase-reversing sinusoidal grating stimuli separated into 5 blocks of 100 phase-reversal pairs, with each block separated by 30 seconds of gray screen (**Figure 1C, D**). Aligning and averaging all stimulus-evoked LFP waveforms occurring within a 400-ms time window from the start of each phase-reversal revealed a stereotyped visually evoked potential (VEP), with an average magnitude that increased across days of exposure (**Figure 1E**). On day 1, the trough-to-peak magnitude of VEPs acquired from control *Grin1*^fl/fl^ mice that had received AAV8-CaMKIIa-GFP virus was significantly lower than for their *Grin1*^fl/fl^ littermates that had received AAV8-CaMKIIa-Cre-GFP to knockout (KO) NMDAR expression in excitatory neurons (**Figure 1F, G**, *median trough-to-peak VEP magnitude difference: -146*.*81 µV, 95% CI = [-221*.*67 -80*.*53] µV, n = 13 Cre*^−^ *control mice, 14 Cre*^+^ *KO mice*). Thus, baseline differences in response magnitude must be considered when comparing data from the control and KO groups.

We next sought to determine how postnatal deletion of functional NMDARs in V1 excitatory cells influences SRP. We induced SRP using a standard protocol, exposing mice to a phase-reversing stimulus at the same orientation each day for 6 consecutive days. We used non-parametric hierarchical bootstrapping for both visualization and statistical evaluation of the data. Briefly, we conducted random draws (with replacement) from the animals and trials that comprised a group, calculated a statistic (such as VEP magnitude) from the average data, then repeated this process 1000 times to generate a distribution. Histograms from this bootstrap were created to aid visualization of the statistical comparison being made (**Figure 1G**). Consistent with previous studies (Frenkel et al., 2006; Cooke et al., 2015; Kaplan et al., 2016; Fong et al., 2020; Hayden et al., 2021), VEP magnitudes increased over days in control animals (**Figure 1F, G**, top panel) such that the VEP on day (d) 6 is significantly larger than d1 (**Figure 1G**, top panel, *median trough-to-peak VEP magnitude difference: 222*.*31 µV, 95% CI = [159*.*46 282*.*45] µV, n = 13 Cre*^−^ *control mice*). For the KO mice, the magnitude also increased over days (**Figure 1F, G**, bottom panel) and the VEP on d6 was significantly larger than d1 (**Figure 1G**, bottom panel, *median trough-to-peak VEP magnitude difference: 76*.*16 µV, 95% CI = [7*.*28 164*.*58] µV, n = 14 Cre*^+^ *KO mice*), but the difference was clearly smaller than control animals.

To directly compare experience-dependent changes in the VEP magnitude between KO and littermate controls, we calculated a term we call the “disparity”. Again, utilizing non-parametric hierarchical bootstrapping we randomly sampled and calculated the d6 - d1 VEP magnitude difference for control animals and for KO animals. These two values were then compared, and the process was repeated (1000 times), yielding a distribution of disparities. If two groups expressed the same level of experience-dependent response plasticity, then the disparity would be 0 µV. However, if the control group expressed more experience-dependent response plasticity, then the disparity would be shifted towards positive numbers.

As expected from visual inspection of **Figure 1G**, CaMKIIa-NMDAR KO animals had significantly less experience-dependent response plasticity than their littermate controls (**Figure 1H**, *median disparity: 146*.*61 µV, 95% CI = [35*.*69 231*.*58] µV, n = 13 Cre*^−^ *control mice, 14 Cre*^+^ *KO mice*). Combined with the day 1 data, this finding suggests that CaMKIIa-NMDAR KO animals either underwent reduced novelty-induced modulation upon initial exposure to a grating stimulus or less experience-dependent potentiation with stimulus familiarization. Regardless, the effects of experience-dependent plasticity are clearly muted in these KO animals.

A similar disparity was also observed on day 7, when interleaved blocks of familiar and novel stimuli were shown. Control animals exhibited lower magnitude VEPs in response to novel stimuli (**Figure 1I**, top panel), showing a statistically significant difference between the magnitude of VEPs produced by familiar and novel stimuli (**Figure 1J**, top panel, *median trough-to-peak VEP magnitude difference: 216*.*68 µV, 95% CI = [156*.*83 275*.*68] µV, n = 13 Cre*^−^ *control mice*). For the KO mice, while a significant difference was apparent between familiar and novel VEPs (**Figure 1I**, bottom panel), that difference was small (**Figure 1J**, bottom panel, *median trough-to-peak VEP magnitude difference: 61*.*43 µV, 95% CI = [16*.*02 106*.*15] µV, n = 14 Cre*^+^ *KO mice*). A group difference in experience-dependent plasticity is inferred from the disparity between CaMKIIa-NMDAR KO animals and their littermate controls (**Figure 1K**, *median disparity: 153*.*46 µV, 95% CI = [84*.*01 227*.*00] µV, n =13 Cre*^−^ *control mice, 14 Cre*^+^ *KO mice*). Thus, NMDARs in CaMKIIa-expressing excitatory principal neurons for required for the full induction and/or expression of SRP.

### Deletion of NMDAR in L6 cortico-thalamic neurons disrupts SRP

NMDAR expressed in excitatory neurons within L4 are not necessary for full SRP expression (Fong et al., 2020). Considered with the above data, we conclude that excitatory cells in other layers must be responsible for the NMDAR-dependence of SRP. Thus, we ablated NMDAR from a population of neurons in another thalamo-recipient layer, L6 (see **Figure 2Ai**), and tested for SRP deficits measured via the L4 VEP. We used a genetic strategy to create a L6 NMDAR knockout mouse line by crossing the *Grin1*^fl/fl^ mice with Neurotensin Receptor 1 (Ntsr1)-Cre mice (see **Methods**), which express Cre within a population of excitatory neurons in L6 that project back to thalamus and have a characteristic pattern of intracortical connectivity (Gong et al., 2007). We confirmed that these mice lack NMDAR in L6 by whole cell recording from pyramidal excitatory neurons (**Figure 2Aii**). GFP-expressing L6 pyramidal neurons from Ntsr1-Cre NMDAR KO mice had a significantly reduced NMDA/AMPA ratio compared to control cells (**Figure 2B, C**, *all bootstrapped 99% confidence intervals between control groups and L6 KO cells did not include 0*), indicating successful reduction in functional NMDARs.

As with the CaMKIIa-NMDAR KO animals, on day 1 the VEP magnitude for the Ntsr1-NMDAR KO mice was increased compared to Cre^−/-^ littermate controls (**Figure 2D**, *median trough-to-peak VEP magnitude difference: -77*.*80 µV, 95% CI = [-144*.*19 -18*.*17] µV, n = 12 control mice, 15 KO mice*). Thus, as with the CaMKIIa-NMDAR KO animals, a similar baseline offset must be considered when comparing data from the Ntsr1-NMDAR KO and the control population. We then induced SRP over 6 days and found that the VEP of both KO and control groups increased over days (**Figure 2D, E**) but, in another phenocopy of the CaMKIIa-NMDAR KO mice, the Ntsr1-NMDAR KO underwent notably less potentiation from day 1 onwards. For the littermate control group, the VEP magnitude on day 6 was significantly greater than day 1 (**Figure 2E**, top panel, *median trough-to-peak VEP magnitude difference: 206*.*79 µV, 95% CI = [140*.*89 276*.*15] µV, n = 12 control mice*). For the KO mice, there was a smaller but still significant VEP difference between day 6 and day 1 (**Figure 2E**, bottom panel, *median trough-to-peak VEP magnitude difference: 60*.*61 µV, 95% CI = [2*.*27 115*.*15] µV, n = 15 KO mice*). As expected from visual inspection of **Figure 2E**, Ntsr1-NMDAR KO animals have less experience-dependent plasticity than their littermate controls (**Figure 2F**, *median disparity: 148*.*59 µV, 95% CI = [65*.*06 239*.*78] µV, n = 12 control mice, 15 KO mice*). Combined with the day 1 data (**Figure 2D**) and like CaMKIIa-NMDAR KO animals, this finding suggests that Ntsr1-NMDAR KO animals either fail to potentiate with stimulus familiarity or fail to suppress cortical response when presented with a novel stimulus on day 1.

As expected, this group disparity is also seen in the familiar/novel difference (**Figure 2G**). Control animals have lower magnitude VEPs in response to novel stimuli (**Figure 2H**, top panel, *median trough-to-peak VEP magnitude difference: 214*.*55 µV, 95% CI = [153*.*54 264*.*31] µV, n = 12 control mice*). For the KO mice, while there was a slight difference between VEPs produced by familiar and novel stimuli (**Figure 2G**), that difference was not significant (**Figure 2H**, bottom panel, *95% CI includes 0, median trough-to-peak VEP magnitude difference: 53*.*20 µV, 95% CI = [-0*.*63 115*.*29] µV, n = 15 KO mice*). This reduction in experience-dependent plasticity is revealed by the disparity between Ntsr1-NMDAR KO animals and their littermate controls (**Figure 2I**, *median disparity: 158*.*17 µV, 95% CI = [73*.*23 237*.*33] µV, n = 12 control mice, 15 KO mice*). Thus, NMDARs are required in Ntsr1-expressing L6 principal cells for the full expression of SRP.

### Experience-dependent changes in the VEP and CSD are seen at multiple depths

To date, measurement of SRP has been restricted to L4 VEPs. The current finding that SRP is disrupted by deletion of NMDAR in L6 principal neurons highlights the paucity of data available for this form of experience-dependent plasticity in layers outside L4. Thus, we obtained acute, laminar recordings of V1 from awake, head-fixed, wild-type C57BL/6J mice to measure SRP across all layers simultaneously. After two consecutive days of habituation, mice were presented with 10 blocks of 100 1-Hz phase-reversals of an oriented grating stimulus. Preceding each block was a period of gray screen, then black, then gray (**Figure 3A**). The gray screens were isoluminant with the phase-reversing grating to minimize pupil dilation artifacts at the onset and offset of stimulus blocks. The black screen was shown for spectral normalization because gray screens elicit luminance-dependent narrow-band oscillations at 60 Hz that are measured in the cortex but arise from subcortical sources (Saleem et al., 2017). This viewing protocol was shown for 4 consecutive days, and on day 5 we recorded acutely from V1 (see **Methods** for recording timeline) while presenting blocks of both familiar and novel stimuli in a pseudo-randomly interleaved manner.

**Figure 3 –.**
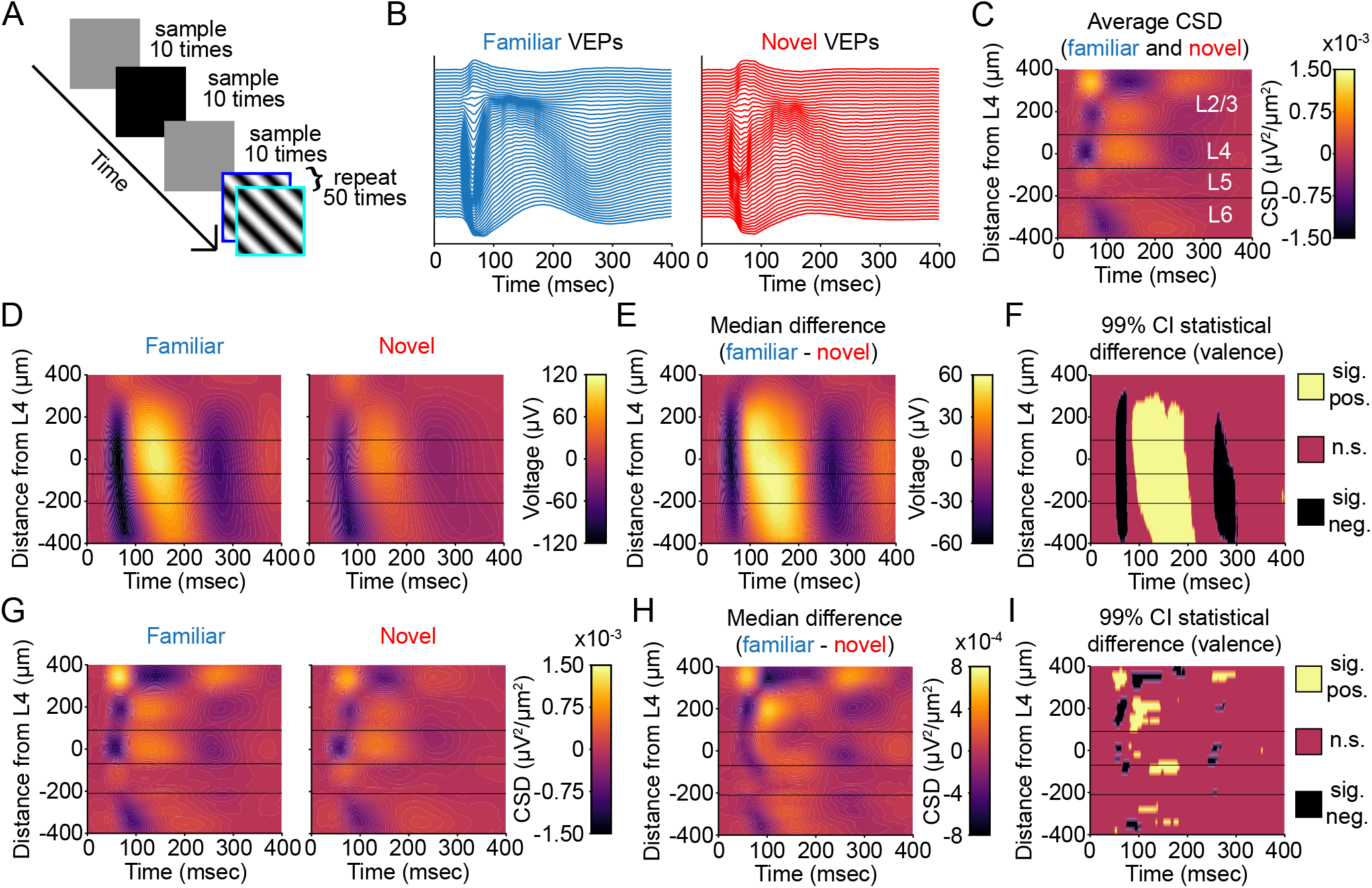
SRP of VEPs across multiple layers of V1. (**A**) We recorded extracellular activity from primary visual cortex (V1) in awake, head-fixed mice in response to phase-reversing sinusoidal grating stimuli (N = 12 mice). The experimental paradigm shows both gray and black screen stimuli between blocks of phase-reversing gratings, with the former ensuring isoluminance at the onset of high-contrast stimuli and the latter used for spectral normalization (see **Methods**). Cortical responses elicited by both the 0° (“flip”, blue) and 180° (“flop”, cyan) phases of stimulation are combined in subsequent analyses. (**B**) Laminar probes were inserted perpendicular to the cortical surface of binocular V1 and recorded at 25 kHz. Plots display the low-pass local field potential from each site of the linear electrode array distributed along the Y-axis according to laminar depth. Aligning to phase-reversal onset reveals stereotyped evoked potentials for familiar and novel stimuli. (**C**) The average current-source density plot for all phase-reversals for familiar and novel stimuli reveals L4 as the site of the earliest and deepest sink. (**D**) The average VEPs across all layers during familiar and novel stimulus blocks. (**E**) Non-parametric hierarchical bootstrapping results for median VEP differences across layers. (**F**) Regions of the laminar VEP in (**E**) where the bootstrapped 99% confidence interval does not include 0 (thus the difference is statistically significant). (**G, H, I**) As in (**D, E, F**), but for the current-source density analysis of the VEPs. All plots use an average smoothing kernel that spans 10% of each axis. Three horizontal lines arranged from top to bottom represent the approximate boundaries between L2/3, L4, L5, and L6..

On the recording day, we obtained VEPs across all layers for both familiar and novel stimuli (**Figure 3B**). Performing current-source density (CSD) analysis on the VEPs across all layers allowed us to confirm the identity of L4, which is the earliest and deepest sink below the superficial source (**Figure 3C**) (Mitzdorf, 1985, 1987; Aizenman et al., 1996). Characteristic VEP morphology observed at this depth further confirmed the location. Visual inspection of the CSD suggested layer boundaries denoted by horizontal lines on laminar plots. These separate the data visually into L2/3, L4, L5, and L6, with statistics reported referencing to these laminar boundaries.

Previous work studying SRP has revealed that long-term stimulus familiarity increases the VEP magnitude in L4, increases the low-frequency (∼15 Hz) power, and decreases the high-frequency (∼65 Hz) power of the LFP recorded within L4 (Frenkel et al., 2006; Cooke and Bear, 2010; Cooke et al., 2015; Kaplan et al., 2016; Fong et al., 2020; Kim et al., 2020; Hayden et al., 2021). We wondered to what extent these changes are exclusive to L4. Plotting the VEP for both familiar and novel stimuli, as well as the difference between them, revealed that visual experience changes the VEP magnitude across most cortical lamina (**Figure 3D-F**). In agreement with previous results, L4 shows a significantly different VEP peak-negativity to peak-positivity difference between familiar and novel stimuli (**Figure 3E, F**, *L4: peak depth = 20 µm, median trough-to-peak difference = 155*.*75 µV, 99% CI = [96*.*74 214*.*86] µV, n = 12 mice*). This difference is also detectable in L2/3 (**Figure 3E, F**, *L2/3: peak depth = 100 µm, median trough-to-peak difference = 146*.*83 µV, 99% CI = [88*.*22 208*.*29] µV, n = 12 mice*), in L5 (**Figure 3E, F**, *L5: peak depth = -80 µm, median trough-to-peak difference = 149*.*30 µV, 99% CI = [96*.*48 205*.*23] µV, n = 12 mice*), and in L6 (**Figure 3E, F**, *L6: peak depth = -220 µm, median trough-to-peak difference = 116*.*29 µV, 99% CI = [74*.*24 170*.*58] µV, n = 12 mice*). Thus, experience-dependent plasticity can be detected in superficial, middle, and deep layers of V1 using the VEP.

The local field potential is sensitive to volume conduction, so the measured VEP magnitude in one layer could be a consequence of activity in other layers. To isolate layer-specific changes, we utilized the CSD, wherein sinks represent flow of electrical current into cells. As expected, L4 is the site of the earliest and deepest sink for both familiar and novel stimuli (**Figure 3G**). All layers display a deeper sink to familiar stimuli compared to novel stimuli. The earliest sink differences appear in L4 (**Figure 3H, I**, *L4: depth = 20 µm, time = 62 msec, sink difference = -7*.*07*10*^*-4*^ *µV*^*2*^*/µm*^*2*^, *99% CI = [-1*.*39*10*^*-3*^ *-1*.*73*10*^*-4*^*] µV*^*2*^*/µm*^*2*^, *n = 12 mice*). Sink differences also appear in L2/3 (**Figure 3H, I**, *L2/3: depth = 340 µm, time = 98 msec, sink difference = -1*.*57*10*^*-3*^ *µV*^*2*^*/µm*^*2*^, *99% CI = [-3*.*48*10*^*-3*^ *-3*.*16*10*^*-4*^*] µV*^*2*^*/µm*^*2*^, *n = 12 mice*), L5 (**Figure 3H, I**, *L5: depth = -80 µm, time = 75 msec, sink difference = -7*.*00*10*^*-4*^ *µV*^*2*^*/µm*^*2*^, *99% CI = [-1*.*35*10*^*-3*^ *-1*.*40*10*^*-4*^*] µV*^*2*^*/µm*^*2*^, *n = 12 mice*), and deep L6 (**Figure 3H, I**, *L6: depth = -360 µm, time = 84 msec, sink difference = -6*.*74*10*^*-4*^ *µV*^*2*^*/µm*^*2*^, *99% CI = [-1*.*13*10*^*-3*^ *- 2*.*43*10*^*-4*^*] µV*^*2*^*/µm*^*2*^, *n = 12 mice*). Thus, experience-dependent plasticity can be detected in superficial, middle, and deep layers of V1 using the CSD, indicating multiple potential sites of synaptic modification.

### Experience-dependent changes in the spectrum of LFP oscillations are seen at multiple depths

Oscillatory analysis of the continuous LFP signal can reveal consistent periodic structure in the electrical activity that reflects changing engagement of local neural circuitry. Because the portion of the recording containing the VEP violates second-order stationarity required for oscillatory analysis, we focused our analysis on the last 600 ms of each 1000 ms presentation when this violation no longer occurs. This approach is consistent with previous work (Chalk et al., 2010; Zhou et al., 2016; Hayden et al., 2021). Additionally, and in line with our previous work (Hayden et al., 2021), we normalized the raw spectrogram to the median spectrogram generated during black screen presentation. This normalized spectrum has units of decibel (dB).

We investigated whether the frequency composition of the LFP changes as a result of visual experience, as we have previously reported (Hayden et al., 2021). The normalized spectra during familiar and novel stimulus viewing showed the two expected findings: novel stimuli elicited more high-frequency power and familiar stimuli elicited more low-frequency power (**Figure 4A**). The experience-dependent change in low-frequency power could be seen in all layers. The largest difference was in L4 (**Figure 4B, C**, *L4: depth = -60 µm, peak frequency = 10*.*01 Hz, median difference = 2*.*33 dB, 99% CI = [1*.*30 3*.*39] dB, n = 12 mice*). However, L2/3 (**Figure 4B, C**, *L2/3: depth = 100 µm, peak frequency = 11*.*23 Hz, median difference = 1*.*99 dB, 99% CI = [1*.*06 3*.*05] dB, n = 12 mice*), L5 (**Figure 4B, C**, *L5: depth = -80 µm, peak frequency = 10*.*01 Hz, median difference = 2*.*29 dB, 99% CI = [1*.*28 3*.*36] dB, n = 12 mice*), and L6 (**Figure 4B, C**, *L6: depth = -220 µm, peak frequency = 10*.*01 Hz, median difference = 1*.*93 dB, 99% CI = [1*.*08 2*.*81] dB, n = 12 mice*) also display this change. The experience-dependent change in high-frequency power could also be seen in most layers. The largest difference was evident in L2/3 (**Figure 4B, C**, *L2/3: depth = 100 µm, peak frequency = 57*.*13 Hz, median difference = -1*.*25 dB, 99% CI = [-1*.*94 -0*.*66] dB, n = 12 mice*), but robust modulation was also observed in L4 (**Figure 4B, C**, *L4: depth = 80 µm, peak frequency = 57*.*13 Hz, median difference = -1*.*24 dB, 99% CI = [-1*.*92 -0*.*63] dB, n = 12 mice*). In L5 the power difference was dampened (**Figure 4B, C**, *L5: depth = -80 µm, peak frequency = 57*.*37 Hz, median difference = -0*.*97 dB, 99% CI = [-1*.*55 -0*.*43] dB, n = 12 mice*), and in L6 it was evident only toward the superficial boundary (**Figure 4B, C**, *L6: depth = -220 µm, peak frequency = 57*.*37 Hz, median difference = -0*.*47 dB, 99% CI = [-0*.*90 -0*.*03] dB, n = 12 mice*). Thus, experience changes oscillatory activity in all layers, with modulation of low and high frequencies most prominent in superficial and middle layers, respectively.

**Figure 4 –.**
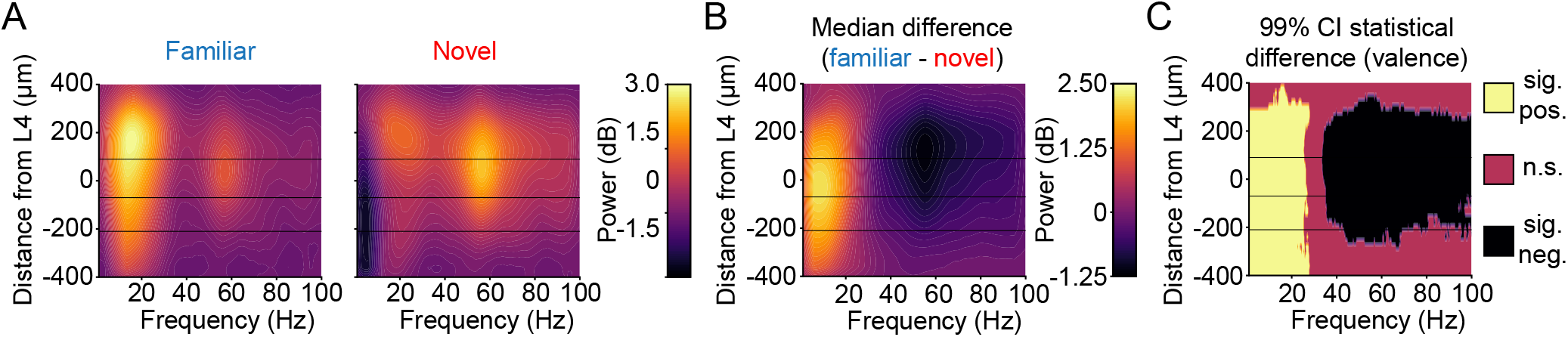
The normalized oscillatory power spectrum across multiple layers of V1 during familiar and novel stimulus viewing. (**A**) The laminar spectrum (normalized to black screen) during familiar and novel stimulus blocks (N = 12 mice). (**B**) Non-parametric hierarchical bootstrapping results for median normalized spectrum differences across layers. (**C**) Regions of the laminar VEP in (**B**) where bootstrapped 99% confidence interval does not include 0 (thus the difference is statistically significant). All plots use an average smoothing kernel that spans 10% of each axis. Three horizontal lines from top to bottom represent the boundaries between L2/3, L4, L5, and L6 as determined by inspection of the CSD.

### Familiarity decreases the average multiunit activity of superficial, middle, and deep layers

To investigate the spiking activity of V1, we calculated a value known as multiunit activity envelope (MUAe, **Figure 5A, B**). MUAe provides an instantaneous measure of the number and size of action potentials of neurons in the vicinity of the electrode tip (Legatt et al., 1980; Brosch et al., 1995; Super and Roelfsema, 2005). This allows us to use every channel in a recording, even if single units are not clearly isolatable, and it does not rely on an arbitrary threshold-crossing to assess multiunit activity. The raw MUAe has a large increase in deeper layers (**Figure 5C**), likely corresponding to increased spiking activity in L5 (Senzai et al., 2019).

**Figure 5 –.**
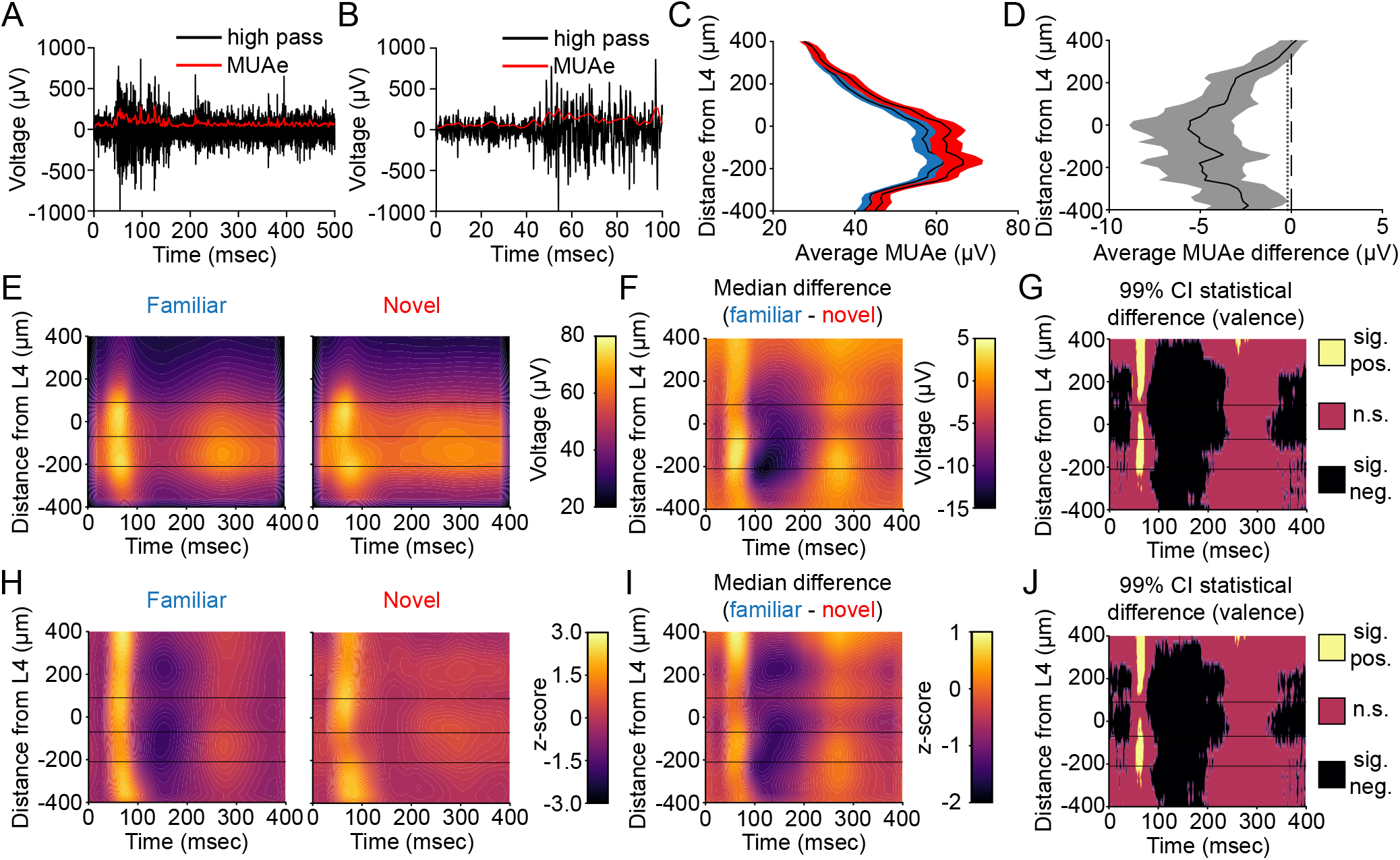
SRP expressed in the multiunit activity across multiple layers of V1. (**A-B**) We calculated the envelope of multi-unit activity (MUAe, see **Methods**) across layers of V1 (N = 12 mice). Example images show that MUAe closely tracks the raw high-pass filtered data. (**C**) The average evoked MUAe during familiar and novel stimulus blocks (collapsed from the first 400 ms of each phase reversal). Averaged activity is presented as mean ± SEM. (**D**) Non-parametric hierarchical bootstrapping results for average MUAe activity across layers. Data are presented as median bootstrap bounded by the 99% confidence intervals. Marks to the left of the vertical dashed 0 line indicate layers with a statistically significant F vs N difference (i.e. where the 99% bootstrapped difference does not contain 0). (**E**) The time course and amplitude of the average MUAe following phase reversals during familiar and novel stimulus blocks. (**F**) Non-parametric hierarchical bootstrapping results for median MUAe differences across layers. (**G**) Regions of the laminar MUAe in (**F**) whose bootstrapped 99% confidence interval does not include 0 (thus the difference is statistically significant). (**H, I, J**) As in (**E, F, G**), but for the z-scored (to gray screen) MUAe. All plots use an average smoothing kernel that spans 10% of each axis. Three horizontal lines represent the boundaries between L2/3, L4, L5, and L6 as determined by inspection of the CSD.

Previous studies have reported a decrease in average neural activity to familiar stimuli (Kim et al., 2020; Gao et al., 2021). We indeed find a small (5-10%), but significant, reduction in the average MUAe to familiar stimuli in L2/3 (**Figure 5D**, *L2/3: depth = 100 µm, median difference = -3*.*68 µV, 99% CI = [-5*.*93 -1*.*78] µV, n = 12 mice*), L4 (**Figure 5D**, *L4: depth = - 20 µm, median difference = -5*.*67 µV, 99% CI = [-8*.*70 -3*.*15] µV, n = 12 mice*), L5 (**Figure 5D**, *L5: depth = -80 µm, median difference = -5*.*06 µV, 99% CI = [-8*.*44 -2*.*72] µV, n = 12 mice*), and L6 (**Figure 5D**, *L6: depth = -220 µm, median difference = -4*.*90 µV, 99% CI = [-7*.*06 -2*.*65] µV, n = 12 mice*). Thus, as with previous studies, familiarity reduces the average neural activity in V1.

However, while the *average* neural activity reduces in response to stimulus experience, previous results from our lab have shown an increase in *peak* activity to familiar stimuli (Cooke et al., 2015). Thus, we next looked at the event related MUAe (**Figure 5E**). Visual inspection reveals a stereotyped difference between familiar and novel stimuli across all layers. This pattern featured a short-latency increase in peak activity followed by a relative quiescent period between phase reversals of the familiar stimulus, as compared to the more consistent activity observed during presentation of the novel stimulus (**Figure 5F, G**). To quantify this, we report the difference in average maximum activity (i.e. peak) within the first 100 ms of the phase-reversal. We also report the difference in the minimum activity (i.e. trough) within the subsequent 100 ms. The largest peak difference is in L5 (**Figure 5F, G**, *L5: depth = -160 µm, peak time = 63 msec, median difference = 17*.*62 µV, 99% CI = [2*.*45 45*.*01] µV, n = 12 mice*). Peak differences are also seen in L2/3 (**Figure 5F, G**, *L2/3: depth = 160 µm, peak time = 63 msec, median difference = 9*.*10 µV, 99% CI = [2*.*27 17*.*46] µV, n = 12 mice*), L4 (**Figure 5F, G**, *L4: depth = -60 µm, peak time = 65 msec, median difference = 10*.*34 µV, 99% CI = [4*.*23 16*.*41] µV, n = 12 mice*), and superficial L6 (**Figure 5F, G**, *L6: depth = -220 µm, peak time = 65 msec, median difference = 12*.*84 µV, 99% CI = [2*.*10 22*.*39] µV, n = 12 mice*). Relative to novel, the familiar stimulus elicits a period of quiescence that is most prominent in L5 (**Figure 5F, G**, *L5: depth = -180 µm, trough time = 144 msec, median difference = -17*.*09 µV, 99% CI = [-29*.*52 -8*.*54] µV, n = 12 mice*) and superficial L6 (**Figure 5F, G**, *L6: depth = -240 µm, trough time = 103 msec, median difference = -16*.*84 µV, 99% CI = [-28*.*31 -8*.*82] µV, n = 12 mice*), but is also evident in L2/3 (**Figure 5F, G**, *L2/3: depth = 100 µm, trough time = 136 msec, median difference = -9*.*86 µV, 99% CI = [-14*.*46 -5*.*53] µV, n = 12 mice*) and L4 (**Figure 5F, G**, *L4: depth = -60 µm, trough time = 134 msec, median difference = -11*.*99 µV, 99% CI = [-16*.*81 -7*.*13] µV, n = 12 mice*).

To aid visualization, we z-scored the stimulus-evoked activity of both familiar and novel stimuli to the same average and standard deviation values (see **Methods, Figure 5H**). This allowed us to make direct comparisons between familiar and novel stimuli in all cortical layers while an preserving an optimal range for the color scale used in each plot. These values are unitless. The z-scored MUAe difference for familiar and novel data showed the expected peak activity followed by a quiescent period (**Figure 5I, J**). A z-scored peak difference exists in all layers (**Figure 5I, J**, *L2/3: depth = 400 µm, peak time = 66 msec, median difference = 2*.*31, 99% CI = [1*.*21 3*.*39]; L4: depth = -60 µm, peak time = 65 msec, median difference = 1*.*28, 99% CI = [0*.*47 2*.*05]; L5: depth = -120 µm, peak time = 65 msec, median difference = 1*.*68, 99% CI = [0*.*78 2*.*65]; L6: depth = -220 µm, peak time = 65 msec, median difference = 1*.*46, 99% CI = [2478 2*.*47]; n = 12 mice*). Additionally, a z-scored trough difference exists in all layers (**Figure 5I, J**, *L2/3: depth = 220 µm, trough time = 142 msec, median difference = -1*.*79, 99% CI = [-2*.*71 -0*.*98]; L4: depth = -60 µm, trough time = 134 msec, median difference = - 1*.*59, 99% CI = [-2*.*29 -0*.*89]; L5: depth = -180 µm, trough time = 144 msec, median difference = -1*.*78, 99% CI = [-2*.*87 -0*.*82]; L6: depth = -220 µm, trough time = 136 msec, median difference = -1*.*72, 99% CI = [-2*.*52 -1*.*02]; n = 12 mice*). Thus, long-term stimulus familiarity increases the peak activity in superficial, middle, and deep layers of V1 prior to a period of quiescence that is potentially attributable to the rapid recruitment of strong feedback inhibition.

In **Figure 6** we compare three different measures of learned stimulus familiarity recorded simultaneously in L4. The MUAe shows both the augmented peak firing and the downward “DC shift” in activity between phase reversals (**Figure 6Ai**). To compare the unit data with VEPs, we normalized the MUAe to the pre-phase-reversal baseline (**Figure 6aii**). Normalization clearly shows how the firing rate *change* from baseline is dramatically augmented when familiar stimuli are phase reversed compared to novel. The peak and trough of the multiunit activity closely align with the negative-going and the positive-going component of the VEP, respectively (**Figure 6Aiii**). For such a simple measure, the L4 VEP is remarkably sensitive and information rich when it comes to detecting experience-dependent changes in cortical information processing. Moreover, the same L4 VEP recording electrode detects changes in activity between phase reversals as revealed by oscillatory activity in the LFP (**Figure 6B**).

**Figure 6 –.**
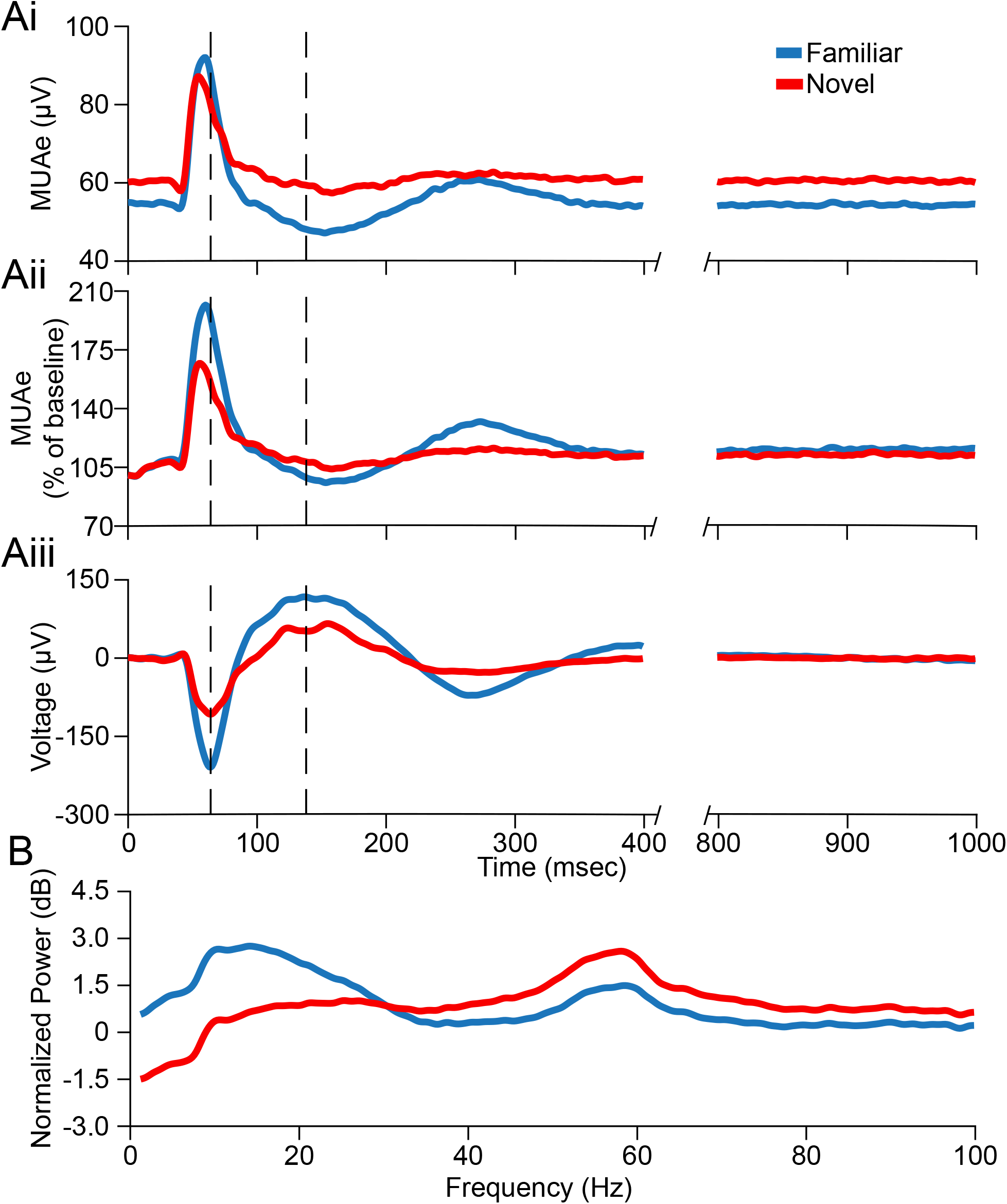
Comparison of SRP expressed in the multiunit activity, VEP, and normalized spectrum in L4 of V1. (**Ai**) The average MUAe during familiar and novel stimulus blocks for L4 (replotted from **Figure 5E**). Familiar stimuli induce lower overall activity, but a phase reversal causes an increase in peak activity compared to novel that is followed by a prolonged period of quiescence. (**Aii**) As in (**Ai**) but normalized to the average pre-phase-reversal baseline. (**Aiii**) The average VEP during familiar and novel stimulus blocks for L4 (replotted from **Figure 3D**). Familiar stimuli induce a larger peak-to-peak VEP magnitude. Dashed vertical lines in **Ai, Aii**, and **Aiii** are the timepoints where the familiar VEP has the maximal negativity and positivity, respectively. (**B**) The average normalized spectrum during familiar and novel stimulus blocks for L4 (replotted from **Figure 4A**). This spectrum is calculated using the local field potential data between 400 and 1000 milliseconds post-phase-reversal-onset to prevent the contamination of the spectral estimate by the VEP response (see **Methods**). Familiar stimuli induce an increase in the low frequency power and a decrease in high frequency power.

## Discussion

Despite abundant evidence supporting the hypothesis that potentiation of thalamocortical synapses is the molecular basis for SRP (Frenkel et al., 2006; Cooke and Bear, 2010), it is now very clear that this does not occur on excitatory neurons in L4 (Montgomery et al., 2021). Indeed, recent findings indicate that SRP expressed as VEP potentiation in L4 is most parsimoniously explained by an experience-dependent reduction in inhibition mediated by PV+ interneurons in this layer (Kaplan et al., 2016; Hayden et al., 2021). Our aims in the current study were therefore to reexamine participation of excitatory neurons in the mechanisms of SRP, with a particular focus on L6 cells known to be a direct target of thalamocortical input, and to perform a laminar analysis of how SRP is expressed across all layers of V1.

To accomplish our first aim, we deleted NMDARs, first from excitatory neurons across all layers, and then from a specific population of L6 projection neurons for which we had genetic access. These *Ntsr1-*expressing L6 neurons were of particular interest, first, because they have been shown to strongly modulate inhibition in L4 through a population of PV+ neurons with translaminar axonal projections, and second, because they project back to the thalamus where they can modulate feedforward activation of V1 (Olsen et al., 2012; Bortone et al., 2014; Kim et al., 2014; Guo et al., 2017; Voigts et al., 2020). With respect to SRP of VEPs in L4, knockout of NMDARs only within *Ntsr1-*expressing L6 neurons phenocopied the loss of NMDARs from excitatory neurons across all layers of V1, supporting the conclusion that these L6 neurons are of particular significance for expression of SRP in L4. This finding is consistent with previous work showing molecular manipulations of L6 neurons in higher order visual cortex strongly influence object recognition memory (Lopez-Aranda et al., 2009), and stands in stark contrast to what was observed when NMDARs were deleted in L4 principal neurons (Fong et al., 2020). Interpretation of the SRP phenotype observed after knocking out NMDARs in L6 is not straight forward, however. Had loss of NMDARs only blocked experience dependent synaptic plasticity in L6, we would expect to observe no baseline change in VEP amplitude as well as no SRP. However, instead we observed a clear increase in VEP magnitude at baseline in response to stimuli the mice had not previously experienced. Thus, the diminished effect of subsequent visual experience could either reflect an impairment in NMDAR-dependent feedforward synaptic plasticity (e.g., impaired LTP (Cooke and Bear, 2010)), a failure to consolidate SRP via corticothalamic feedback (Durkin et al., 2017), or the partial occlusion of SRP caused by reduced activation of these L6 neurons in the absence of NMDARs. More work is needed to parse these possible explanations, but we can confidently conclude based on available data that L6 neurons contribute to a circuit that plays a critical role in the expression of SRP in L4 VEPs.

The profound influence of a molecular manipulation of L6 on plasticity measured in L4 underscored the need to develop a more thorough description of SRP across all layers of V1. This was accomplished using translaminar multichannel electrodes which allowed analysis of VEPs and spiking activity elicited by phase reversals of novel and familiar stimuli, as well as changes in oscillatory activity in the LFP between phase reversals. Across superficial, middle, and deep cortical layers we observed that VEP magnitudes and the spectral content of V1 activity elicited by familiar and novel stimuli were strikingly different, extending previous findings restricted to L4 (Frenkel et al., 2006; Cooke and Bear, 2010; Cooke et al., 2015; Kaplan et al., 2016; Fong et al., 2020; Kim et al., 2020; Hayden et al., 2021). Other than responses measured near the cortical surface, VEP magnitudes were elevated in response to phase reversals of familiar stimuli compared to novel across all layers, notably including L6 (**Figure 3**). CSD analysis revealed the expected translaminar progression of information flow in V1 (Mitzdorf, 1985; Aizenman et al., 1996) and confirmed augmentation of both short- and long-latency current sinks in all layers except deep L5. Similarly, the signature of stimulus familiarity in the LFP oscillations measured between phase reversals was clearly present in all layers (**Figure 4**). However, whereas the enhanced power in the low-frequency (∼15 Hz) band occurred at all depths, reduced power in the high-frequency gamma band (∼65 Hz) appeared to be confined to the middle layers, centered on L4. This finding is of interest, as decreased gamma is associated with reduced recruitment of PV+ interneurons (Chen et al., 2017; Veit et al., 2017), which likely accounts for VEP potentiation in L4 (Kaplan et al., 2016).

We found comparable modulation of multiunit activity in superficial, middle, and deep layers of V1 by familiar visual stimuli that is best described as a temporal redistribution of cell firing (**Figure 5**). In all layers, there was an increase in the multiunit peak firing in response to phase reversals of a familiar stimulus compared to a novel stimulus. However, this averaged phasic increase in firing rate was quickly followed by a prolonged period of quiescence between phase reversals, leading to an overall reduction in population activity during familiar stimulus viewing compared to novel. This finding is consistent with observations using intracellular recordings from single neurons in superficial V1, showing a reduction in average firing rate to sustained viewing of familiar stimuli (Gao et al., 2021). It is of interest to compare our electrophysiological observations with previous studies using two-photon calcium imaging to estimate cellular activity. Imaging of pyramidal cells in superficial layers and L4 have shown consistently a reduction in averaged calcium sensor fluorescence in response to familiar stimuli in comparison to novel stimuli (Kato et al., 2015; Makino and Komiyama, 2015; Kim et al., 2020). The study by Kim et al. (2020) in L4 is particularly relevant, as it utilized the same SRP protocol we used in the current study. We think it is likely that the fleeting increase in firing after each phase reversal, lasting tens of milliseconds, was likely missed due to the relatively poor temporal resolution of the imaging method. In any case, these differences highlight how use of phase reversing stimuli and electrophysiology to probe modifications of V1 that accompany visual learning can yield novel mechanistic insights that could be missed using other stimulation and recording methods (e.g., drifting gratings and calcium imaging). Without getting too speculative about the precise circuit elements involved, our data are generally consistent with a model in which learning occurs by enhancement of net feedforward thalamocortical excitation (revealed by the phasic response potentiation) that, in turn, recruits increased polysynaptic feedback inhibition that quenches the activity between phase reversals. In this model, the reduced tonic activation of cortex during familiar stimulus viewing is a consequence of a net increase in inhibitory tone rather than a long-term depression of excitatory synaptic transmission.

In conclusion, in addition to showing how profoundly stimulus familiarity influences activity across all layers of mouse V1, we have identified a surprising influence of L6 excitatory neurons on producing pronounced differences in the response of L4 to familiar and novel stimuli. Further work will be required to determine if this influence occurs through a PV+ inhibitory intermediary as suggested by our previous observations (Kaplan et al., 2016; Hayden et al., 2021) and, if so, whether these intermediaries reside within L6 (Bortone et al., 2014; Frandolig et al., 2019) or in more superficial layers (Kim et al., 2014). In addition, further work will be required to understand if the influence of *Ntsr1*-expressing L6 neurons occurs via their characteristic feedback to the primary sensory thalamus. The activity of these L6 neurons may control the trade-off between stimulus detection and perception of stimulus features (Guo et al., 2017). Thus, the switch in the mode of activity in canonical cortical circuitry produced by L6 CT neurons potentially serves the primary function of habituation by limiting the influence of familiar sensory experience on attention and energy use, while enhancing vigilance for unexpected changes in the environment.

## Acknowledgements

We acknowledge the invaluable support of Arnold Heynen, Ming-fai Fong, Nina Palisano, Jessica Buckey, Athene Wilson-Glover, Kiki Chu, and Erin Hickey. Feng-Ju Weng, Maia Lee, Julie (Heejung) Kim, and Christian Candler contributed to collection of pilot and control data related to this project. The research was supported by NIH grant R01 EY023037, the Picower Institute Innovation Fund, and the Picower Fellows program. Samuel Cooke would also like to acknowledge current funding from the Biotechnology and Biological Sciences Research Council (BBSRC) grant (BB/S008276/1).

## References

Aizenman CD, Kirkwood A, Bear MF (1996) A current source density analysis of evoked responses in slices of adult rat visual cortex: implications for the regulation of long-term potentiation. Cereb Cortex 6:751–758.

Aton SJ, Suresh A, Broussard C, Frank MG (2014) Sleep promotes cortical response potentiation following visual experience. Sleep 37:1163–1170.

Bokil H, Andrews P, Kulkarni JE, Mehta S, Mitra PP (2010) Chronux: a platform for analyzing neural signals. J Neurosci Methods 192:146–151.

Bortone DS, Olsen SR, Scanziani M (2014) Translaminar inhibitory cells recruited by layer 6 corticothalamic neurons suppress visual cortex. Neuron 82:474–485.

Brosch M, Bauer R, Eckhorn R (1995) Synchronous high-frequency oscillations in cat area 18. Eur J Neurosci 7:86–95.

Chalk M, Herrero JL, Gieselmann MA, Delicato LS, Gotthardt S, Thiele A (2010) Attention reduces stimulus-driven gamma frequency oscillations and spike field coherence in V1. Neuron 66:114–125.

Chen G, Zhang Y, Li X, Zhao X, Ye Q, Lin Y, Tao HW, Rasch MJ, Zhang X (2017) Distinct Inhibitory Circuits Orchestrate Cortical beta and gamma Band Oscillations. Neuron 96:1403–1418 e1406.

Collingridge GL (2003) The induction of N-methyl-D-aspartate receptor-dependent long-term potentiation. Philos Trans R Soc Lond B Biol Sci 358:635–641.

Cooke SF, Bear MF (2010) Visual experience induces long-term potentiation in the primary visual cortex. J Neurosci 30:16304–16313.

Cooke SF, Bear MF (2014) How the mechanisms of long-term synaptic potentiation and depression serve experience-dependent plasticity in primary visual cortex. Philos Trans R Soc Lond B Biol Sci 369:20130284.

Cooke SF, Ramaswami M (2020) Ignoring the innocuous: mechanisms of habituation. In: The Cognitive Neurosciences, 6th Edition (Poeppel D, Mangun GR, Gazzaniga MS, eds), pp 197–206. USA: MIT Press.

Cooke SF, Komorowski RW, Kaplan ES, Gavornik JP, Bear MF (2015) Visual recognition memory, manifested as long-term habituation, requires synaptic plasticity in V1. Nat Neurosci 18:262–271.

Cruikshank SJ, Lewis TJ, Connors BW (2007) Synaptic basis for intense thalamocortical activation of feedforward inhibitory cells in neocortex. Nat Neurosci 10:462–468.

Douglas RJ, Martin KA (2004) Neuronal circuits of the neocortex. Annu Rev Neurosci 27:419–451.

Durkin J, Suresh AK, Colbath J, Broussard C, Wu J, Zochowski M, Aton SJ (2017) Cortically coordinated NREM thalamocortical oscillations play an essential, instructive role in visual system plasticity. Proceedings of the National Academy of Sciences of the United States of America 114:10485–10490.

Fong MF, Finnie PS, Kim T, Thomazeau A, Kaplan ES, Cooke SF, Bear MF (2020) Distinct Laminar Requirements for NMDA Receptors in Experience-Dependent Visual Cortical Plasticity. Cereb Cortex 30:2555–2572.

Frandolig JE, Matney CJ, Lee K, Kim J, Chevee M, Kim SJ, Bickert AA, Brown SP (2019) The Synaptic Organization of Layer 6 Circuits Reveals Inhibition as a Major Output of a Neocortical Sublamina. Cell Rep 28:3131–3143 e3135.

Frenkel MY, Sawtell NB, Diogo AC, Yoon B, Neve RL, Bear MF (2006) Instructive effect of visual experience in mouse visual cortex. Neuron 51:339–349.

Gao M, Lim S, Chubykin AA (2021) Visual Familiarity Induced 5-Hz Oscillations and Improved Orientation and Direction Selectivities in V1. J Neurosci 41:2656–2667.

Gong S, Doughty M, Harbaugh CR, Cummins A, Hatten ME, Heintz N, Gerfen CR (2007) Targeting Cre recombinase to specific neuron populations with bacterial artificial chromosome constructs. J Neurosci 27:9817–9823.

Grill-Spector K, Henson R, Martin A (2006) Repetition and the brain: neural models of stimulus-specific effects. Trends in cognitive sciences 10:14–23.

Guo W, Clause AR, Barth-Maron A, Polley DB (2017) A Corticothalamic Circuit for Dynamic Switching between Feature Detection and Discrimination. Neuron 95:180–194 e185.

Hayden DJ, Montgomery DP, Cooke SF, Bear MF (2021) Visual recognition is heralded by shifts in local field potential oscillations and inhibitory networks in primary visual cortex. J Neurosci.

Kaneko M, Fu Y, Stryker MP (2017) Locomotion Induces Stimulus-Specific Response Enhancement in Adult Visual Cortex. J Neurosci 37:3532–3543.

Kaplan ES, Cooke SF, Komorowski RW, Chubykin AA, Thomazeau A, Khibnik LA, Gavornik JP, Bear MF (2016) Contrasting roles for parvalbumin-expressing inhibitory neurons in two forms of adult visual cortical plasticity. Elife 5.

Kato HK, Gillet SN, Isaacson JS (2015) Flexible Sensory Representations in Auditory Cortex Driven by Behavioral Relevance. Neuron 88:1027–1039.

Kim J, Matney CJ, Blankenship A, Hestrin S, Brown SP (2014) Layer 6 corticothalamic neurons activate a cortical output layer, layer 5a. J Neurosci 34:9656–9664.

Kim T, Chaloner FA, Cooke SF, Harnett MT, Bear MF (2020) Opposing Somatic and Dendritic Expression of Stimulus-Selective Response Plasticity in Mouse Primary Visual Cortex. Frontiers in cellular neuroscience 13:555.

Kirkwood A, Bear MF (1994) Hebbian synapses in visual cortex. J Neurosci 14:1634–1645.

Legatt AD, Arezzo J, Vaughan HG, Jr. (1980) Averaged multiple unit activity as an estimate of phasic changes in local neuronal activity: effects of volume-conducted potentials. J Neurosci Methods 2:203–217.

Lopez-Aranda MF, Lopez-Tellez JF, Navarro-Lobato I, Masmudi-Martin M, Gutierrez A, Khan ZU (2009) Role of layer 6 of V2 visual cortex in object-recognition memory. Science 325:87–89.

Makino H, Komiyama T (2015) Learning enhances the relative impact of top-down processing in the visual cortex. Nat Neurosci 18:1116–1122.

Mitzdorf U (1985) Current source-density method and application in cat cerebral cortex: investigation of evoked potentials and EEG phenomena. Physiol Rev 65:37–100.

Mitzdorf U (1987) Properties of the evoked potential generators: current source-density analysis of visually evoked potentials in the cat cortex. Int J Neurosci 33:33–59.

Montgomery DP, Hayden DJ, Chaloner FA, Cooke SF, Bear MF (2021) Stimulus-Selective Response Plasticity in Primary Visual Cortex: Progress and Puzzles. Front Neural Circuits 15:815554.

Olsen SR, Bortone DS, Adesnik H, Scanziani M (2012) Gain control by layer six in cortical circuits of vision. Nature 483:47–52.

Poort J, Khan AG, Pachitariu M, Nemri A, Orsolic I, Krupic J, Bauza M, Sahani M, Keller GB, Mrsic-Flogel TD, Hofer SB (2015) Learning Enhances Sensory and Multiple Non-sensory Representations in Primary Visual Cortex. Neuron 86:1478–1490.

Rankin CH, Abrams T, Barry RJ, Bhatnagar S, Clayton DF, Colombo J, Coppola G, Geyer MA, Glanzman DL, Marsland S, McSweeney FK, Wilson DA, Wu CF, Thompson RF (2009) Habituation revisited: an updated and revised description of the behavioral characteristics of habituation. Neurobiology of learning and memory 92:135–138.

Saleem AB, Lien AD, Krumin M, Haider B, Roson MR, Ayaz A, Reinhold K, Busse L, Carandini M, Harris KD (2017) Subcortical Source and Modulation of the Narrowband Gamma Oscillation in Mouse Visual Cortex. Neuron 93:315–322.

Saravanan V, Berman GJ, Sober SJ (2020) Application of the Hierarchical Bootstrap to Multi-Level Data in Neuroscience. bioRxiv:819334.

Senzai Y, Fernandez-Ruiz A, Buzsaki G (2019) Layer-Specific Physiological Features and Interlaminar Interactions in the Primary Visual Cortex of the Mouse. Neuron 101:500–513 e505.

Speed A, Del Rosario J, Burgess CP, Haider B (2019) Cortical State Fluctuations across Layers of V1 during Visual Spatial Perception. Cell Rep 26:2868–2874 e2863.

Super H, Roelfsema PR (2005) Chronic multiunit recordings in behaving animals: advantages and limitations. Progress in brain research 147:263–282.

Thompson RF, Spencer WA (1966) Habituation: a model phenomenon for the study of neuronal substrates of behavior. Psychol Rev 73:16–43.

Tsien JZ, Chen DF, Gerber D, Tom C, Mercer EH, Anderson DJ, Mayford M, Kandel ER, Tonegawa S (1996) Subregion- and cell type-restricted gene knockout in mouse brain. Cell 87:1317–1326.

Ulbert I, Halgren E, Heit G, Karmos G (2001) Multiple microelectrode-recording system for human intracortical applications. J Neurosci Methods 106:69–79.

Veit J, Hakim R, Jadi MP, Sejnowski TJ, Adesnik H (2017) Cortical gamma band synchronization through somatostatin interneurons. Nat Neurosci 20:951–959.

Voigts J, Deister CA, Moore CI (2020) Layer 6 ensembles can selectively regulate the behavioral impact and layer-specific representation of sensory deviants. Elife 9.

Zhou H, Schafer RJ, Desimone R (2016) Pulvinar-Cortex Interactions in Vision and Attention. Neuron 89:209–220.

